# ElectroPen: An ultralow-cost piezoelectric electroporator

**DOI:** 10.1101/448977

**Authors:** Gaurav Byagathvalli, Soham Sinha, Yan Zhang, Mark P. Styczynski, Janet Standeven, M. Saad Bhamla

## Abstract

Electroporation is a basic yet powerful method for delivering small molecules (RNA, DNA, drugs) across cell membranes by application of an electrical field. Due to its vital role, electroporation has wide applicability from genetically engineering cells, to drug- and DNA-based vaccine delivery. Despite its broad applications in biological research, the high cost of electroporators is an obstacle for many budget-conscious laboratories. To address this need, we describe a simple, inexpensive, and hand-held electroporator inspired by a common household piezoelectric gas lighter. The proposed “ElectroPen” device costs about 20 cents, is portable (13 g), is fabricated on-demand using 3D-printing, and delivers repeatable exponentially decaying pulses of about 2000 V in 5 ms. We provide a proof-of-concept demonstration by genetically transforming plasmids into *E. coli* strains and show comparable transformation efficiency and cell growth with commercial devices, but at a fraction of the cost. Our results are validated by an independent team across the globe, providing a real-world example of democratizing science through frugal tools. Thus, the simplicity, accessibility, and affordability of our device holds potential for making modern synthetic biology accessible in high-school, community, and field-ecology laboratories.

## INTRODUCTION

Electroporators are utilized across research laboratories for a wide spectrum of purposes in molecular biology, biotechnology, and biomedical engineering fields [1, 2]. Examples of these applications include bacterial transformation [3], genetic engineering with CRISPR [4], gene transfer in mammalian embryos [5], cancer treatments using electrochemotherapy [6], transdermal drug-delivery [7], and gene-based vaccine-delivery [8]. Due to their versatility, electroporators are in high demand across research laboratories and industries. However, commercial electroporators are complex, expensive, and bulky hardware that can cost upwards of thousands of dollars [9]. In budget-restrained laboratories such as U.S. public high schools, biological field stations in remote places (field-biology), and research laboratories in resource-poor countries, lack of access to electroporators is a significant bottleneck for conducting basic biological research. The need for accessible and affordable electroporators has prompted researchers to develop simple electronic circuits using relays and capacitors [10–14]. However, these devices still cost hundreds of dollars and require extensive electronics and hardware skills to construct, making them impractical. Thus, there is a need for a simple, easy-to-build, and low-cost electroporator for biological research.

We describe the design and implementation of an ultra-low-cost (23 cents (Supplementary Tables 1 and 2), lightweight (13g) (Supplementary Fig. S8), and 3D-printed electroporator inspired by a common household piezoelectric lighter (Fig. 1a, c), and henceforth referred to as the ‘ElectroPen’. We demonstrate that the ElectroPen delivers repeatable, exponentially-decaying electrical pulses with an average of about 2000 V in 5 ms, through action of a high-speed hammer (8 ms^-1^, 30, 000 ms^-2^) that is actuated by mechanical springs and latches, and easily triggered by a human finger. To enable bacterial transformations, we develop inexpensive cuvettes constructed from glass slides and aluminum tape using a facile technique, eliminating the need for expensive commercial cuvettes (Fig. 1b). Using this setup, we demonstrate that the ElectroPen successfully transforms a plasmid encoding constitutive expression of the the Green Fluorescent Protein (GFP) into BL-21 *E. coli* and achieves similar transformation efficiencies as compared to standard electroporators, opening up potential for broad genetic transformation using this low-cost instrument (Supplementary Fig. S1).

**FIG. 1.**
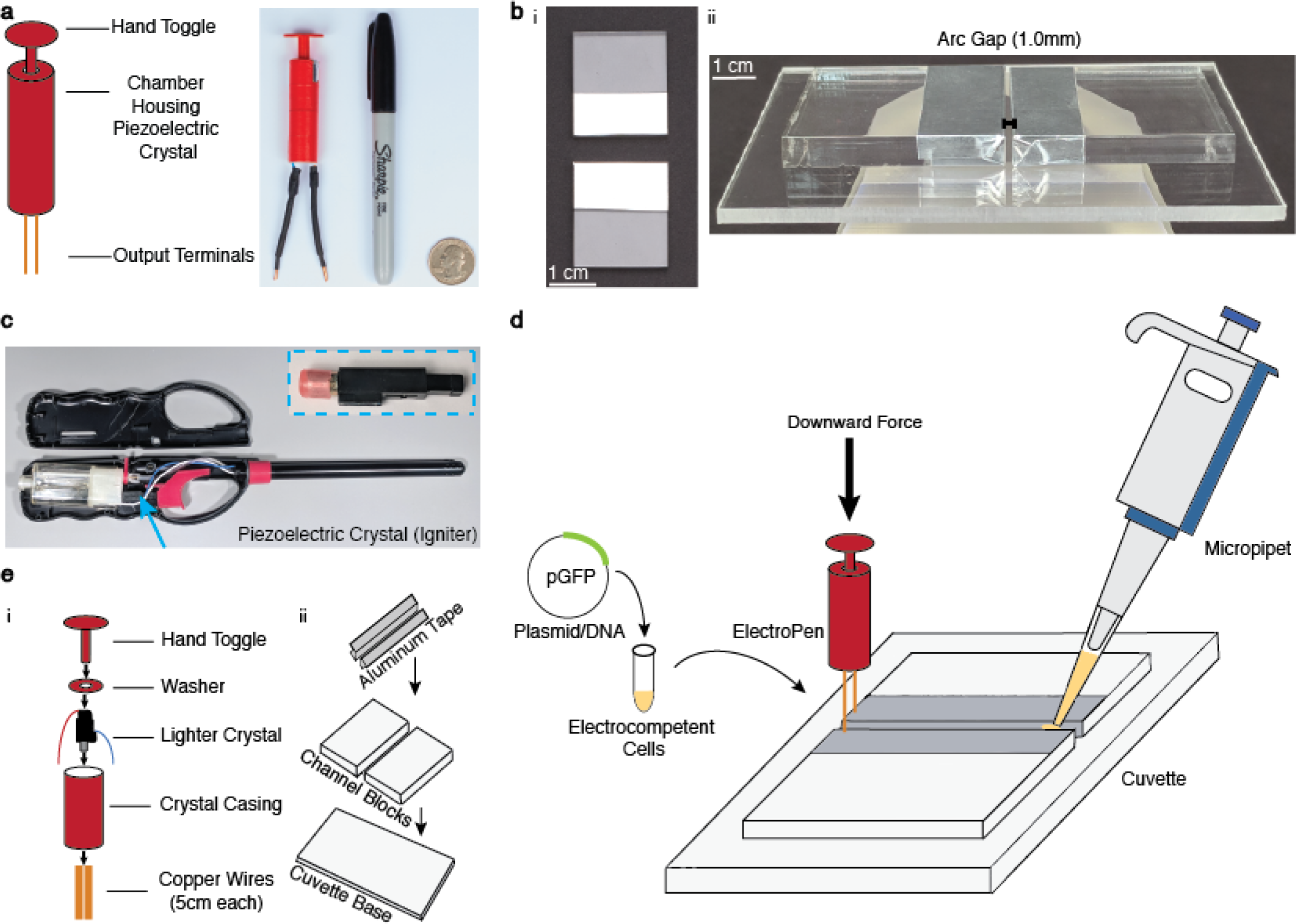
ElectroPen Platform. **a** Design of the 3D-printed low-cost electroporation device along with a depiction of its size scale, demonstrating portability. Device incorporates a simple operation by pressing down to trigger the piezoelectric mechanism, resulting in electrical discharge. **b** Design of the alternative electroporation cuvette. Cuvette design consists of two blocks (shown here in acrylic) covered with aluminum tape to act as electrodes, and placed on a base with a gap distance of 1.0 mm. The cuvette can be built out of other materials (Supplementary Fig. S9) as an alternative for industrial equivalents. **c** Depiction of the origin of the piezoelectric ignition mechanism found within the common stove lighter. It is located next to the butane tank, and the toggle on a lighter directly exerts a force on this mechanism to produce a spark. **d** Illustration of the general protocol for using the ElectroPen system. The cellular suspension is added to the gap in the cuvette, after which the ElectroPen is connected and pressed to trigger a voltage potential. The cell suspension is then recovered in Luria Bertani broth and plated. See Supplementary Movie S4 for a detailed demonstration. **e** Illustration of the individual components of the 3D-printed ElectroPen platform and custom cuvette.

## RESULTS

### ElectroPen design and fabrication

The design of the ElectroPen includes a 3D-printed cylindrical chamber that houses a piezoelectric crystal harvested from a commercial lighter (Fig. 1a, c). The chamber has wire pass-throughs at the bottom and a hand toggle inserted at the top which when pressed downwards provides the equivalent force utilized in a conventional lighter (Fig. 1d, e, and Supplementary Movie S4). The output voltage remains consistent independent of the user’s force, and is discussed in detail later. The simplicity of the design provides for easy construction as an ElectroPen can be fabricated in 15 minutes (Supplementary Movie S2 and Fig. S6). The total cost of materials is 23 cents (Supplementary Table 1).

Due to the high cost of standard cuvettes ($4.54 for 0.1 mm electroporation cuvette by ThermoFisher Scientific P41050) that often have to be purchased in bulk through commercial vendors, we developed a custom cuvette that can be rapidly fabricated with plastic (for example, acrylic) and custom gap-widths can be generated to accommodate different fluid volumes (Fig. 1b, Supplementary Fig. S7, S9). The general design of the cuvette includes a base and two blocks with aluminum tape covering the sides of each block to function as electrodes, and the space in between to hold the competent cell mixture to be transformed (Fig. 1e, Supplementary Fig. S7, Materials and Methods for details). Although we use both a commercial laser cutter setup and a glass slide design, we show that this technique can be easily extended to other materials such as wood, and reliable arc spacing of 1.0 mm can be achieved by using sheets of paper or a credit card to set the gap distance (Supplementary Fig. S9 and Movie S2).

### Exponentially decaying pulses

We next measure the electrical response of the ElectroPen using a high-voltage probe connected to an oscilloscope. The average curves (of n=39 firings from 3 users) follow an exponential function described by V(t) = *V*_0_*e*^(*-αt*)^, where function V is the voltage in kV, t is the time in seconds, *V*_0_ = 1.834 kV is the initial maximum voltage, and *α* = 213.1 s^-1^ is the the exponential decay time to reach one-third of the initial value as shown in Fig. 2a. The exponential decay is a function of the piezoelectric effect occurring through the polarization of the lead zirconate titanate (PZT) crystal (Supplementary Fig. S10). We note that the electrical output is remarkably reproducible over many trials despite being conducted by multiple users due to the spring-based design of the ElectroPen, details of which are discussed in the next section. We optimized the design of the ElectroPen to produce a maximum voltage of V_max_ = 2±0.3 kV with an average time constant *τ* = 5.1±0.9 ms. These values correspond to a field strength of 20 kV/cm, which is optimal for *E. coli* transformation[15]. For a piezoelectric crystal, the theoretical V_*th*_ can be predicted from the following equation:

**FIG. 2.**
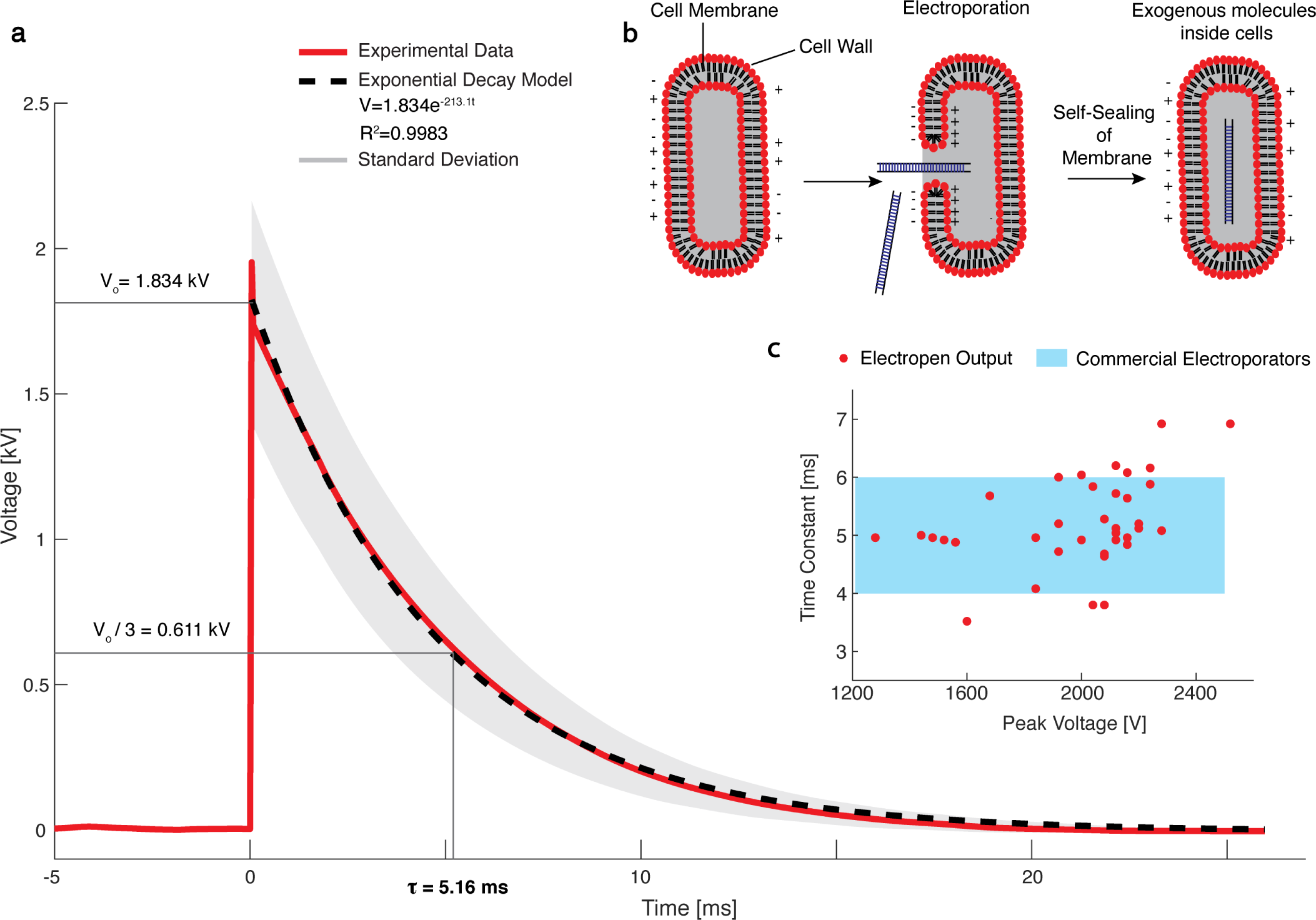
High-voltage output of the ElectroPen for electroporation. **a** Electrical waveform model for the high-voltage pulse produced by the ElectroPen. Piezoelectric output is produced in the form of an exponentially decaying wave (the optimal waveform for electroporation of prokaryotic cells) achieving an average peak voltage output of *V*_max_ = 1997.8 ± 278.2 V and time constant *τ* = 5.1 ± 0.9 ms. In this model, V_0_ indicates the average initial value of the waveforms, and time constant is defined as the time taken for the waveform to decay from its peak voltage to 1/3rd of its peak voltage. **b** Illustration of the theoretical intracellular phenomena that occur during electroporation, in which the formation of aqueous pores is triggered by the high-voltage pulse, allowing for uptake of DNA particles, and decay of the voltage pulse supplemented with nutrient media recovery allows for membrane repair, sealing the DNA particle inside the cell for subsequent expression. **c** The maximum voltage and time constant outputs produced by the ElectroPen are within the range of commercial electroporators.

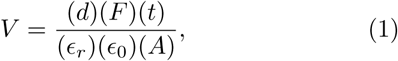

where d = Piezoelectric charge coefficient, F = Force applied on the crystal, t = Thickness of the crystal, *∊*_*r*_ = Relative permittivity, *∊*_0_ = Permittivity of a vacuum, and A = Area of impact surface [16]. For a given ElectroPen design, all the parameters, including force applied on the crystal remain constant. Using published values for the parameters for the PZT crystal (Supplementary Discussion) and a force (F = 10 N), we obtain V_*th*_ ∼ 2.7 kV (Supplementary Discussion), which is of the same order of magnitude as the experimentally measured voltages. Although the current voltages and time constants are designed within the range of values for successful electroporation of *E. coli* as shown in Fig. 2c [15], we demonstrate that the underlying principle of this device (Fig. 2b) can be used to generate V_*max*_ = 30, 000 V (Supplementary Fig. S3), and tuned for a range of time constants and voltage outputs for different biological and biomedical applications[17].

### A latch-spring mechanism for high voltage generation

How does the ElectroPen generate repeatable, consistently high voltages, independent of user force? The ElectroPen exploits a simple and inexpensive mechanical latch-spring mechanism to release a small hammer-pin structure onto the piezoelectric crystal to generate high voltages. In contrast, standard electroporators use complex electrical circuits with costly microprocessor-controlled relays to generate similar voltages. To illustrate the underlying mechanics, we recorded the rapid motion of the piezoelectric crystal using a high-speed camera at 1057 frames per second (Supplementary Movie S1). The mechanism consists of two springs, a hammer (metal piece striking the crystal), and the PZT crystal itself connected to a metal conductor (Fig. 3a, b). The hammer action functions in 3 phases: a loading phase, latch-release phase, and a relaxation phase (Fig. 3c-e).

**FIG. 3.**
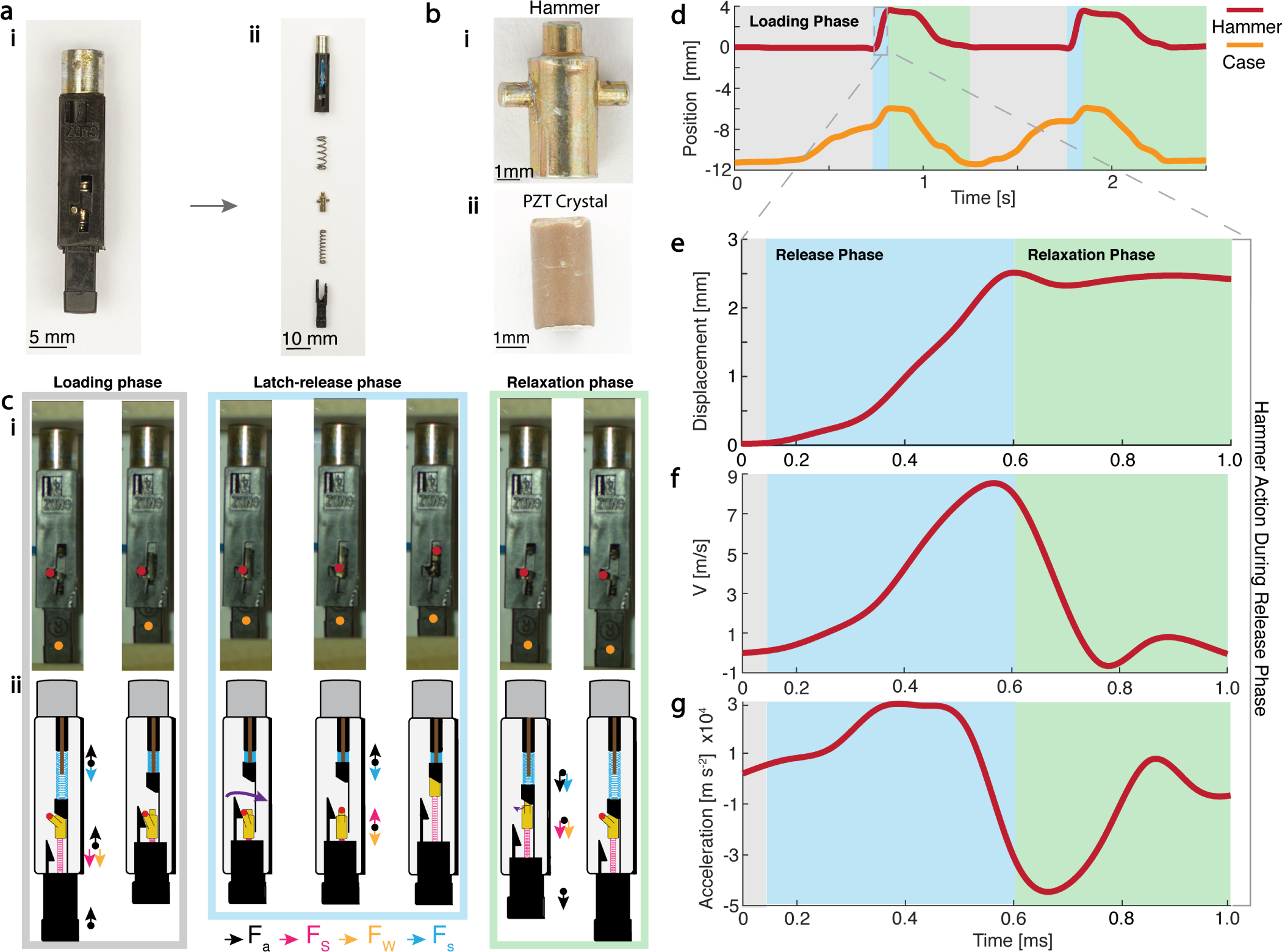
Latch-spring mechanisms for repeatable high-voltage generation. **a** Image of the striking mechanism (hammer action) found within the piezo igniter in a lighter. The parts include from top to bottom: metal conductor housing the piezoelectric crystal, springs, hammer, release-spring, and geometrical latch. The presence of two springs is to decouple the loading and release phase for consistent voltage output. **b** Images of the hammer and lead zirconate titanate (PZT) crystal. The circular surface area of the hammer comes into direct contact with a pin that strikes the piezoelectric crystal, generating a voltage through the piezoelectric effect. **c** (i) Snapshots from high-speed video illustrating the position of the hammer during the loading, latch-release, and relaxation phases. (ii) Free-body diagram indicating movement of each part through each phase of the hammer action, including activation and deactivation of spring forces. **d** Plot of displacement of the hammer and the lower case as a function of time obtained using high-speed image video. **e-g** Zooming into the dynamics of the hammer during the latch-release phase reveals that the hammer achieves a peak velocity of 8 ms^-1^ in 0.5 ms, which corresponds to a remarkable acceleration of 30,000 ms^-2^. The explosive acceleration results in a 10 N force (mass of hammer is 0.3 g) exerted over a tiny area of the PZT crystal.

During the loading phase, the hammer is held in a locked position by a mechanical latch, as the lower spring and upper spring are being compressed. This phase generates spring potential energy by compressing both springs through the user exerted force on the 3D-printed hand toggle. As the compression continues, the wedge-shaped piece of the casing forces the pin of the hammer to rotate outwards of the latch. Once the pin of the hammer has reached a critical point in its rotation, i.e., it is no longer held in place by the latch, the lower spring instantaneously decompresses. This act converts the stored spring energy into the kinetic energy of the hammer, resulting in a high impact force on the cylindrical face of the PZT crystal to generate voltage. Since the degree of lower spring compression is dependent only on the spring-latch design, the quantity of force striking the crystal by the hammer is independent of the user-applied force on the toggle. As a result, the output voltages are remarkably consistent. In the relaxation phase, the user releases the user-applied force to reset the hammer to its initial position.

Analysis of high-speed videos of the hammer releasing indicate that the hammer is able to reach a maximum velocity of 8 m/s at a peak acceleration of almost 30, 000 ms^-2^ or force of 3000 g-force (Fig. 3e-g). Through the explosive nature of the hammer action’s acceleration, a powerful resultant impulsive force of 10 N strikes the PZT crystal, resulting in a high-voltage pulse.

### *E. coli* transformation using ElectroPen

To demonstrate the utility of the ElectroPen in enabling bacterial electrotransformations, we prepared BL-21 *E. coli* electrocompetent cells and transformed them with a recombinant plasmid with constitutive SFGFP expression under the control of J23119 promoter (see Materials and Methods, Supplementary Fig. S12). These plasmids were electroporated into BL-21 electrocompetent *E. coli* using both the ElectroPen with a custom 0.1 cm cuvette and a commercial electroporator and 0.1 cm cuvette (BioRad MicroPulser). By measuring the GFP fluorescence levels using a plate reader, we confirm that the plasmid encoding GFP was successfully electroporated using the ElectroPen (Fig. 4a, Supplementary Fig. S4) in comparison to the negative control (water transformed into BL-21 *E. coli*). The fluorescence intensity values for the ElectroPen trials are within the range of outputs produced by the standard electroporator, indicating successful electroporation, uptake of DNA, and expression of GFP by the *E. coli* bacteria. Moreover, the transformation efficiencies between the ElectroPen and the standard electroporator are within an order of magnitude (Fig. 4b).

**FIG. 4.**
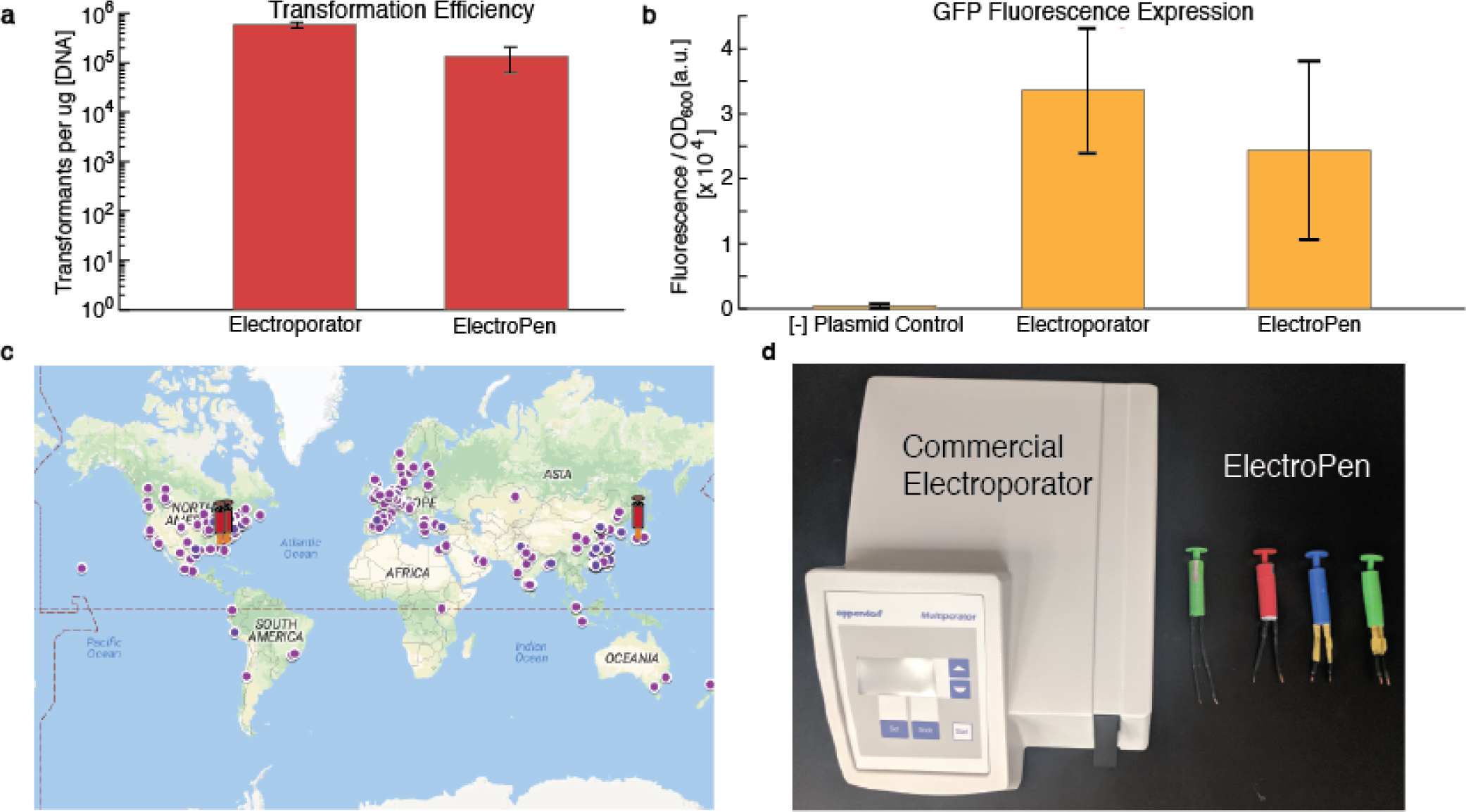
E.coli transformation and distribution of platform globally. **a** The values for the commercial electroporator and ElectroPen are 5.87 ± 0.74 × 10^5^ and 1.35 ± 0.72 × 10^5^ transformants per microgram of DNA, respectively. Thus, the transformation efficiency between a commercial device and ElectroPen are within an order of magnitude. Error Bars, S.D. n = 4 for electroporator and n = 7 for ElectroPen. **b** Plot of fluorescence output from GFP incorporation into *E. coli*. Values from the ElectroPen transformation were compared with a commercial electroporator (BioRad MicroPulser), confirming successful transformation and GFP expression. Here, the negative control is water (no plasmid) transformed into *E.coli*, and electroporator/ElectroPen refers to a plasmid encoding GFP transformed into *E. coli*. Error Bars, S.D. n = 3. **c** Map illustrating the distribution of 2018 iGEM Teams (high school and undergraduate) across the world, with the ElectroPen currently distributed to groups in Georgia, United States and Taipei, Taiwan. **d** Comparison between the commercial electroporator and different ElectroPens.

Lastly, we highlight the rapid dissemination and use of the ElectroPen for synthetic biology experiments through collaboration with two teams from the International Genetically Engineered Machine (iGEM) competition. We shared the device design files, sample protocols and digital instructions with the University of Georgia and Taipei American School (Taiwan) iGEM teams (Fig. 4c, d). These teams of high school students and undergraduates tested the ElectroPen by transforming plasmids encoding GFP *E. coli* into two different strains, DH5a (UGA) DH5*α*/Nissle 1917 (TAS Taipei). The teams obtained successful fluorescence expression and comparable transformation efficiency data, validating the reproducibility and rapid dissemination of the ElectroPen (Supplementary Fig. S2, S4, and S12).

## DISCUSSION

Electroporators are versatile tools for genetic engineering and basic biology. A new push towards frugal science has inspired the development of several devices, such as the FoldScope[18] and PaperFuge[19], which allow expansion of these disciplines into high-schools, under-funded laboratories, and even field research. These low cost devices serve as alternatives for expensive lab equipment, while simultaneously removing numerous barriers including but not limited to cost, access to electricity, and portability. Here, we have theoretically modeled and experimentally validated the functionality and effectiveness of the ElectroPen, the cheapest electroporator in the world (23 cents). We highlight the mechanism of repeatable high-voltage generation through a mechanical spring-latch system in conjunction with a piezoelectric crystal. We also successfully demonstrate application of the first usage of piezoelectric voltage discharges for the electroporation of *E. coli* through the development of the ElectroPen. Through collaboration with national and international iGEM teams, we demonstrate the reproducibility of the ElectroPen and also show that, due to its simplicity, ease of access, and 3D-printed design, the ElectroPen can be easily incorporated in high-school, research, and even citizen-science based community laboratories, enabling modern synthetic biology in budget-constrained environments. We envision the application of the ElectroPen beyond synthetic biology into novel drug and vaccine delivery platforms for low-resource settings in developing nations. Ultimately, the ElectroPen is another example in frugal science that serves to bypass economical and infrastructure limitations in the advancement of scientific research by the next-generation of young scientists across the globe.

## MATERIALS AND METHODS

### ElectroPen System

The mechanism utilized for the production of the voltage output has been obtained from a conventional stove lighter gun. The hammer action mechanism present for the ignition of butane gas has been extracted from the lighter and placed into a 3D-printed casing to allow for more ease of operation and stability. A toggle was added to assist in the exertion of a force on the hammer action that strikes the crystal, and copper wires were attached for conductance and extension of the length of the terminals. Additionally, a 0.1 cm cuvette was designed to serve as a less expensive alternative for standard electroporation cuvettes. It consists of two pieces of acrylic with aluminum tape on the ends to create a gap distance for the electrical arc. This is surrounded by another piece of acrylic to hold the structure in place and create a channel for the cell solution to be held. Other materials can be easily substituted in place of acrylic or glass for fabrication. When the ElectroPen is connected to the cuvette, a spark from the voltage output jumps between the two electrodes, indicating that an electric potential has been established. When cells are placed in the channel, the output voltage travels through the cell solution between the electrodes, allowing electroporation to occur.

### Recombinant Plasmid

The plasmid (pADS001) used in in-house electroporation experiment is a medium copy plasmid (p15A origin) with chloramphenicol resistance. pADS001 constitutively expresses super folder green fluorescent protein (SFGFP) under a strong J23119 promoter and a strong ribosomal binding site. The plasmid is purified with Omega EZNA miniprep kit according to manufacturer’s protocol and sequence confirmed.

### Electrocompetent Cell Preparation

BL-21 *E. coli* cells were inoculated into 5 mL overnight liquid cultures and stored in an 37*°* Celsius incubator set to shake at 170 rpm. After 18 hours of growth, they were diluted to a ratio of 1:100 in sterile Luria Bertani (LB) media and shaken in an 37*°* Celsius incubator at 170 rpm. During this growth phase with the cell culture shaking, the centrifuge was set to 4*°* Celsius. Autoclaved water and 10% glycerol solution were then stored in a refrigerator at 4*°* Celsius in preparation for the following wash steps. The Optical Density (OD) of the cells was continuously monitored every 30 minutes until the cells reached an OD of approximately 0.6. The cells were then transferred into 50 mL conical tubes on ice and allowed to cool for 10 minutes. The suspension was centrifuged at 2500 x g for 6 minutes, after which the supernatant was discarded. The cell pellets were combined by serially resuspending in 13 mL of chilled autoclaved water to concentrate the cell suspensions into two 50 mL conical flasks. The suspension was centrifuged at 2500 x g for 6 minutes, after which the supernatant was discarded. The previous wash step was then repeated. The cell pellets were then resuspended in 13 mL of chilled 10% glycerol solution. The suspension was centrifuged at 2500 x g for 6 minutes, after which the supernatant was discarded. The previous wash step was then repeated. The cell pellets were then resuspended and combined in 4 mL of chilled 10% glycerol and aliquoted into chilled microcentrifuge tubes with 100 *µ*L per tube. They were then stored in a −80*°* Celsius freezer for long-term preservation.

### Electroporation

Cuvettes were stored in a –20*°* Celsius freezer for at least 24 hours before electroporation. The custom cuvette was then taken out of the freezer and sterilized with ethanol before electroporation. During this time, the cuvette was kept on ice. The ElectroPen was then tested on the empty cuvette to ensure that the spark successfully traveled between the electrodes. The frozen DH5a or BL-21 electrocompetent cells thawed on ice for approximately 10 minutes, after which they were aliquoted in 50 *µ*L amounts to individual ice-chilled tubes. 1 *µ*L (2 ng DNA) of the plasmid construct was then added to each cell suspension and tubes were gently shaken to ensure thorough mixing while water was added to a different cell suspension as the negative control. The cell suspension was then transferred into the electrode gap (cuvettes were quickly taken office to a smooth surface for electroporation, but then placed back on ice right after), and shocked once with the ElectroPen. An oscilloscope was connected to the electrodes on the cuvette to determine successful generation of the voltage. Immediately following the shock, 100 *µ*L of pre-warmed (37*°* Celsius) Luria Bertani broth was carefully added to the gap to recover the cell suspension. The recovered suspension was then transferred into a microcentrifuge tube, after which an additional 900 *µ*L of Luria Bertani broth was added to the tube for recovery. The tubes were stored at 37*°* Celsius in an incubator set to shake and allowed to recover for 60-90 minutes. The tubes were centrifuged at 3000 x g for 1 minute, and 800 *µ*L of the supernatant was discarded. The cell pellet was then reconstituted in the remaining LB media and plated onto the appropriate antibiotic plates.

### Efficiency Calculations

Transformation efficiency was calculated using the following variables: a = Number of colonies, b = Volume of transformation solution, c = Volume of DNA added, d = Volume Plated, and e = Concentration of DNA with all volumes in *µL* and DNA mass in *µg*. The following formula was used to calculate transformation efficiency:

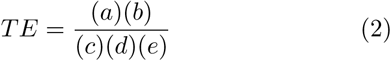

### Fluorescence Measurements

For in vivo fluorescence measurements, three colonies were picked from each of the test cases no plasmid control, plasmid introduced by electroporator, and plasmid introduced by electropen. The colonies from no plasmid controls were inoculated into 5 mL of LB media without antibiotic resistance, and the colonies bearing pADS001 plasmid were inoculated into 5 mL of LB containing chloramphenicol. The cultures were grown at 37*°* and 170 rpm for 18 hrs before 50 *µ*L of the overnight culture were transferred into 950 *µ*L of M9 minimal media (1xM9 minimal salts, 0.01% thiamine hydrochloride, 20 mM Dextrose, 2 mg/mL SC-Ura, 2 mM MgSO_4_, 0.1 mM CaCl_2_) containing chloramphenicol antibiotics. Overnight cultures of no plasmid controls were transfer into M9 without antibiotics. The M9 subcultures were then grown for 4 hours at 37*°* and 180 rpm. After 4 hours, 100 *µ*L of each subculture was transferred onto a clear bottom 96 well plate (Corning 3651) and diluted with 100 *µ*L of 1xPBS. Optimal density (*OD*_600_ nm) and fluorescence (485 nm excitation and 528 nm emission, Gain 60) were measured on a Biotek Synergy H4 microplate reader at room temperature. All experiments were repeated independently three times for a minimum of nine biological replicates per test case.

### Voltage Measurements

A high voltage probe (Elditest CT4026) with a 1000:1 divider ratio was used with an oscilloscope (Teledyne Lecroy WaveAce2014) with a measurement span of 25 ms, and with a vertical scale of 1 kV to measure the output voltage from the ElectroPen (Supplementary Figure S5). The high voltage probe has a grounding wire which was connected to the oscilloscope.

### Replicate Samples

For fluorescence measurements, samples were taken in biological replicates. Colonies from each plate for each trial were isolated and grown in individual liquid cultures, and this was done for three separate trials. The trial average is a combination of the replicates for each trial and average for the sample is the average of the trials for that sample (negative control, electroporator, and ElectroPen). The standard deviation is representative of the variation in data from independent trials.

## ACKNOWLEDGEMENTS

We thank all members of the Bhamla Lab for their feedback. We thank L. Graber and C. Park for assistance in measuring the voltage outputs; H. Shekhani for discussions on piezoelectric theory; C. Chang, L. Tsai, and J. Clapper from Taipei American School iGEM 2018 for their collaboration; S. George and K. McConnell from UGA iGEM 2018 for their collaboration; and Lambert iGEM for their support and collaboration. M.S.B acknowledges funding support through NSF (award no. 181733). Additional information can be found at the BhamlaLab/ElectroPen web page.

## AUTHOR CONTRIBUTIONS

GB, SS, and MSB conceived idea for ElectroPen. GB designed and constructed prototypes. GB and JS conducted biological experiments in *E. coli*. YZ and MPS conducted plate reader measurements and analysis. SS conducted voltage measurements. GB, SS, and MSB con-ducted high-speed experiments. GB, SS, JS, and MSB analyzed the data and wrote the manuscript.

