## Supplementary material for "ElectroPen: An ultralow-cost piezoelectric electroporator": SI Text

### Supplementary Information - ElectroPen: An ultralow-cost piezoelectric electroporator

Gaurav Byagathvalli,[1] Soham Sinha,[2] Yan Zhang,[2] Mark P. Styczynski,[2] Janet Standeven,[1] and M. Saad Bhamla[2,*]
[1]*Lambert High School, 805 Nichols Rd, Suwanee, GA, 30024, USA*
[2]*School of Chemical & Biomolecular Engineering,*
*Georgia Institute of Technology, 311 Ferst Drive NW, Altanta, GA 30332, USA*
(Dated: December 21, 2018)

#### CONTENTS

I. Supplementary Movies 1
A. S1: High-speed dynamics of the Hammer Action 1
B. S2: Construction Process for ElectroPen Device 1
C. S3: Construction Process for the ElectroPen Cuvette 1
D. S4: Protocol for Usage of the ElectroPen 1

II. Supplementary Materials and Methods 2
A. Organization of Experimental Trials 2
B. TAS Taipei Plasmid Construct 2

III. Supplementary Theory Discussion 2
A. Estimation of theoretical voltage for piezoelectric crystal 2

IV. Supplementary Tables 2

V. Supplementary Figures 2

References 2

#### I. SUPPLEMENTARY MOVIES

##### A. S1: High-speed dynamics of the Hammer Action

High-speed video (1057 fps) of the ElectroPen's hammer action proceeding through the described three stages: loading phase, latch-release phase, and relaxation phase. During the loading phase, as the user applies an input force, the casing moves upwards, compressing the lower and upper spring while the hammer remains in the locked position. The applied force results in a wedge pushing the hammer arm out of the latch (locking position). After the hammer arm has completely moved out of the latch, the latch-release phase begins, and the spring extends, projecting the hammer upwards, striking a pin connected to the piezoelectric crystal. The hammer then returns to the original position as the user pulls back on the hammer action restoring the original position.

##### B. S2: Construction Process for ElectroPen Device

Video outlining the overall method for the construction of the ElectroPen. Video speed has been increased, but the average time taken to build an ElectroPen is approximately 15 minutes.

##### C. S3: Construction Process for the ElectroPen Cuvette

Video outlining the overall method for the construction of the ElectroPen cuvette. Video speed has been increased, but the average time taken to build an ElectroPen cuvette is approximately 5 minutes.

##### D. S4: Protocol for Usage of the ElectroPen

Video outlining the overall method for using the ElectroPen. The cuvette for electroporation was kept in a $-20^{\circ}$ freezer for 24 hours prior to electroporation, and cuvette was held on ice prior to conducting the trial.

#### II. SUPPLEMENTARY MATERIALS AND METHODS

##### A. Organization of Experimental Trials

For the data represented in Fig. 4 (a,b), the trials were conducted in subsets. The first group of experiments comprised of one positive control with plasmid introduced using the electroporator, seven experimental samples with plasmid introduced using the ElectroPen, and one negative control with no plasmid. The ElectroPen samples were run first, followed by the electroporator, and then finally the negative control. The following day, 3 samples with the positive control with the electroporator and 4 samples with the negative control with the ElectroPen were conducted. This was to ensure sufficient samples for biological triplicate measurements.

For the data represented in Supplementary Fig. S2, the 3 ElectroPen experimental samples were run first followed by the positive control with the electroporator and negative control with the ElectroPen. This trial was conducted at the University of Georgia with the support of UGA iGEM Team Members in an effort to characterize the ElectroPen's functionality with the DH5a strain of *E. coli*.

##### B. TAS Taipei Plasmid Construct

The plasmid utilized encoded GFP (BBa E0040) under the control of a pLac promoter (BBa R0010) with parts obtained from the iGEM Parts Registry.

#### III. SUPPLEMENTARY THEORY DISCUSSION

##### A. Estimation of theoretical voltage for piezoelectric crystal

The piezoelectric crystal present within the lighter consists of a disc-orientation, with voltage enhanced through longitudinal thickness. The force is applied on the bottom surface, with the thickness (in this case height) of the crystal acting as the amplification factor (Supplementary Fig. S10). The following equations were utilized to calculate the theoretical maximum voltage of the piezoelectric crystal present for the ElectroPen.

$$D_3 = (d_{31})(\sigma_1$$

$D_{31}$ represents the piezoelectric charge constant. We can substitute the variables to include charge, force, and area.

$$\frac{Q}{A_3} = (d_{31})(\frac{F_1}{A_1})$$

We can then substitute charge for capacitance and voltage, and rearrange to include $A_3$ on other other side of the equation.

$$(C_3)(V_4) = (d_{31})(F_1)(\frac{A_3}{A_1})$$

Divide equation by capacitance $= \frac{\epsilon_0 \epsilon_r A_3}{t}$.

$$v_3 = \frac{(d_{31})(F_1)(A_3)(t)}{(\epsilon_0)(\epsilon_r)(A_3)(A_1)}$$

Simplify equation to remove $A_3$ from equation.

$$V_3 = \frac{(d_{31})(F_1)(t)}{(\epsilon_0)(\epsilon_r)(A_1)}$$

Substitute with $g_{33}$ (piezoelectric voltage constant) equation value.

$$\frac{d_{33}}{(\epsilon_0)(\epsilon_r)} = g_{33}$$

$$V_3 = \frac{(g_{33})(F_1)(t)}{A_1}$$

Using values of $g_{33}$ = 0.0265Vm/N [1], $F_1$ = 10 N, t = 8 mm, and A = $\pi * r^2$ where r = 1 mm, we obtain a maximum theoretical output of 2,699.3 V, which is of the same order of magnitude as experimental voltages. The small mis-match in the experimental and theoretical values is attributed to the resistance in the copper wires as well as confinement-effects of the crystal within the plastic case.

#### IV. SUPPLEMENTARY TABLES

#### V. SUPPLEMENTARY FIGURES

[1] K. Harikrishnan, D. V. Bavbande, D. Mohan, B. Manoharan, M. R. Prasad, and G. Kalyanakrishnan, in *IOP Conference Series: Materials Science and Engineering* (2018).

| Part | Cost (in US Dollars) |
|---|---|
| Piezoelectric Crystal | $0.02 |
| Copper Plated Wire | $0.10 |
| Heat-Shrinking Wire Insulator | $0.05 |
| 3D-Printed Casing | $0.05 |
| Aluminum Tape | $0.01 |
| **ElectroPen** | **$0.23** |

TABLE I. List of parts and costs for construction of the ElectroPen. The net price is reflective of goods purchased in wholesale from suppliers as listed above (excluding production costs). Links to the parts: Piezoelectric Crystal, Copper Plated Wire, Heat-Shrinking Wire Insulator, 3D-Printed Casing, and Aluminum Tape.

| Device | Additional Supplies | Cost† | Electricity | Weight |
|---|---|---|---|---|
| BioRad Micropulser Electroporator | Commercial cuvettes* | $2,399.00 | Yes | 2.9kg |
| Eppendorf Eporator | Commercial cuvettes* | $2,619.00 | Yes | 3.2kg |
| BioRad Gene Pulse Cell Microbial Electroporator | Commercial cuvettes* | $6,875.00 | Yes | 6.6kg |
| **ElectroPen** | **Low-cost, custom cuvettes** | **$0.23** | **No** | **13g** |

TABLE II. Comparison between standard commercial electroporators and the ElectroPen. The listed electroporators reflect common equipment utilized in labs. The ElectroPen reflects a fraction of the cost of its industrial equivalent, while not requiring access to electricity and weighing magnitudes less.*0.1/0.2mm Gap Industrial Electroporation Cuvettes. † cost includes only the device.

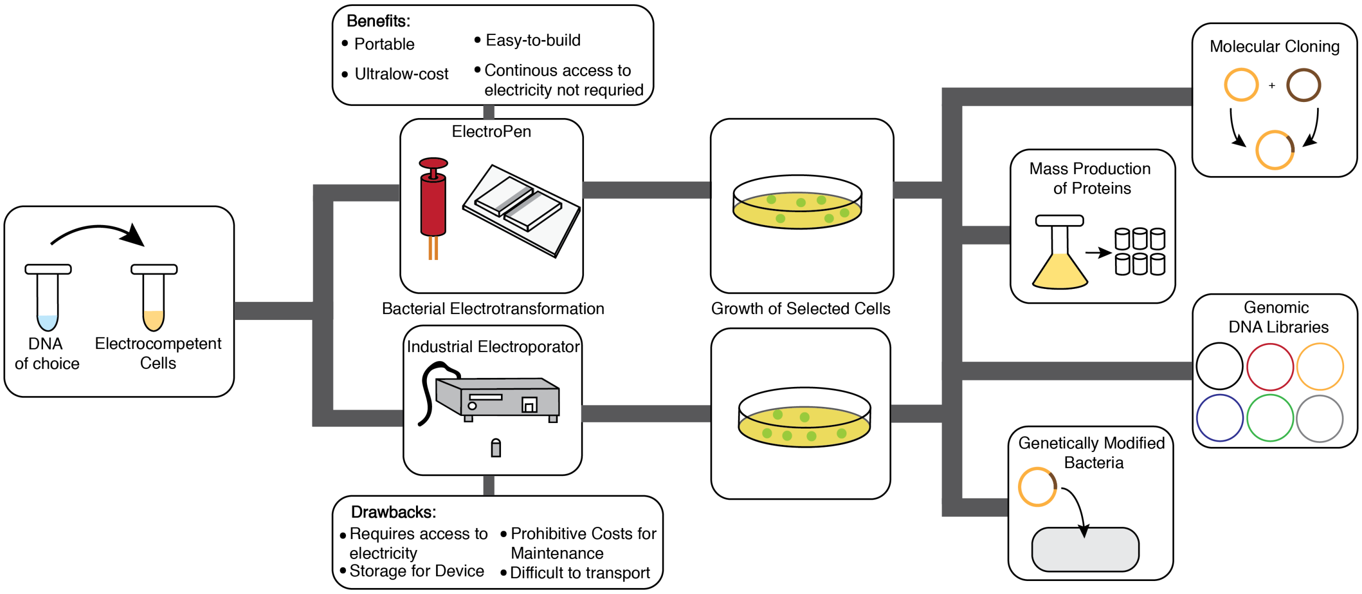


FIG. S1. Workflow schematic for usage and applications of the ElectroPen in comparison to commercial electroporators, as well as their advantages/drawbacks. DNA is added to thawed electrocompetent cells and electroporated using either system and plated on antibiotic plates to isolate successful clones. In terms of applications within bacteria, electroporation can be utilized for molecular cloning, genetic engineering and recombination, mass production, etc., demonstrating the importance of electroporators in these fields. The ElectroPen can be applied for the same purposes without the restrictions of costs, electricity, and portability, demonstrating its potential as a complete alternative for a standard commercial electroporator.

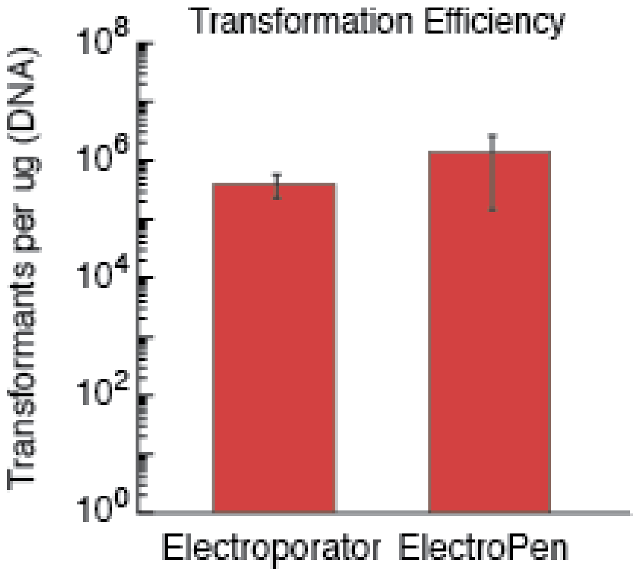


FIG. S2. Transformation efficiency data from the trial conducted at the University of Georgia using DH5a *E. coli*.

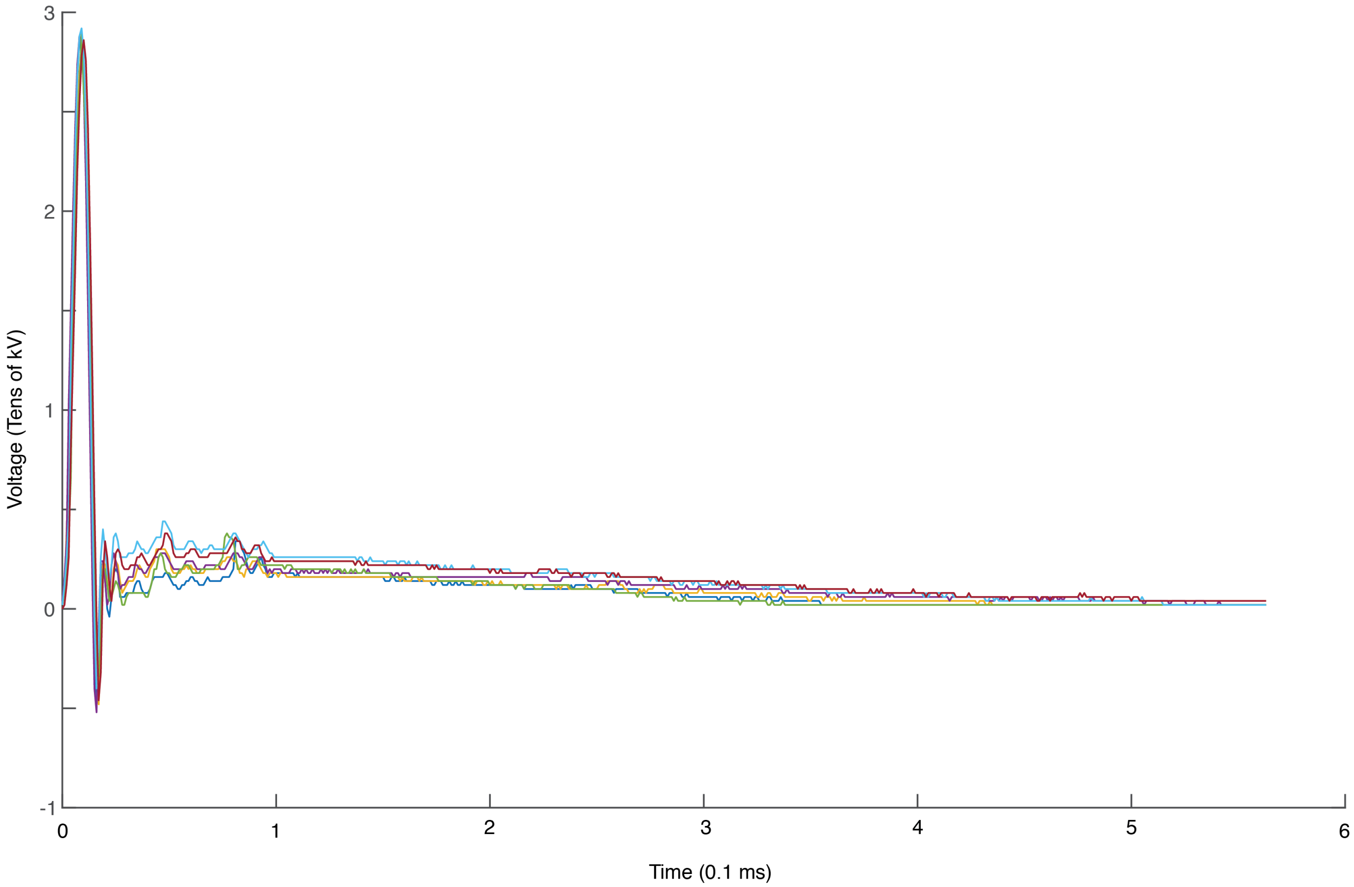


FIG. S3. Waveforms produced by ElectroPen using different piezoelectric crystals demonstrating maximum voltage of 30,000 Volts

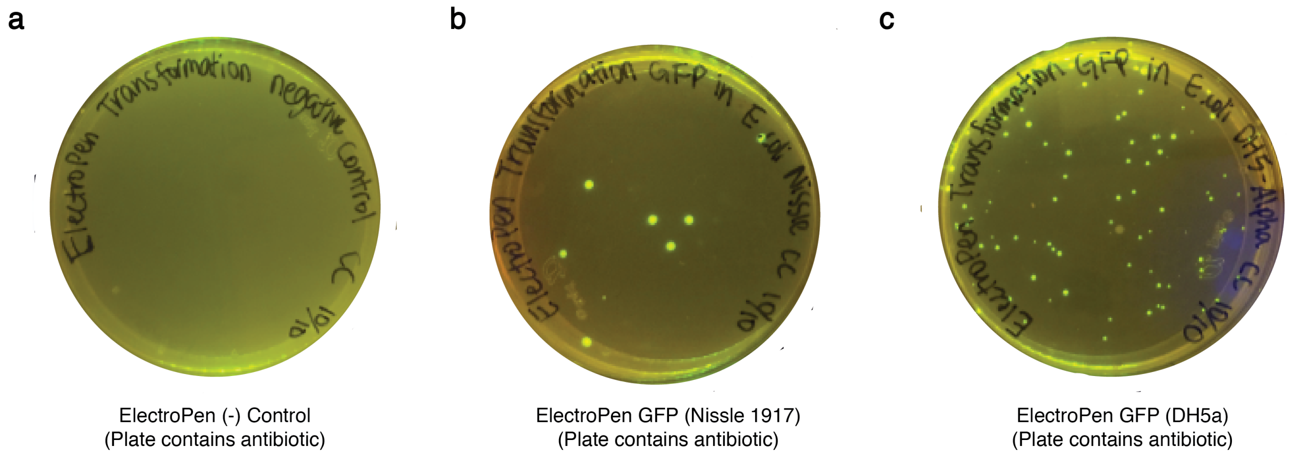


FIG. S4. Images of the plates with fluorescence from the independent collaboration trials conducted by TAS Taipei iGEM 2018.

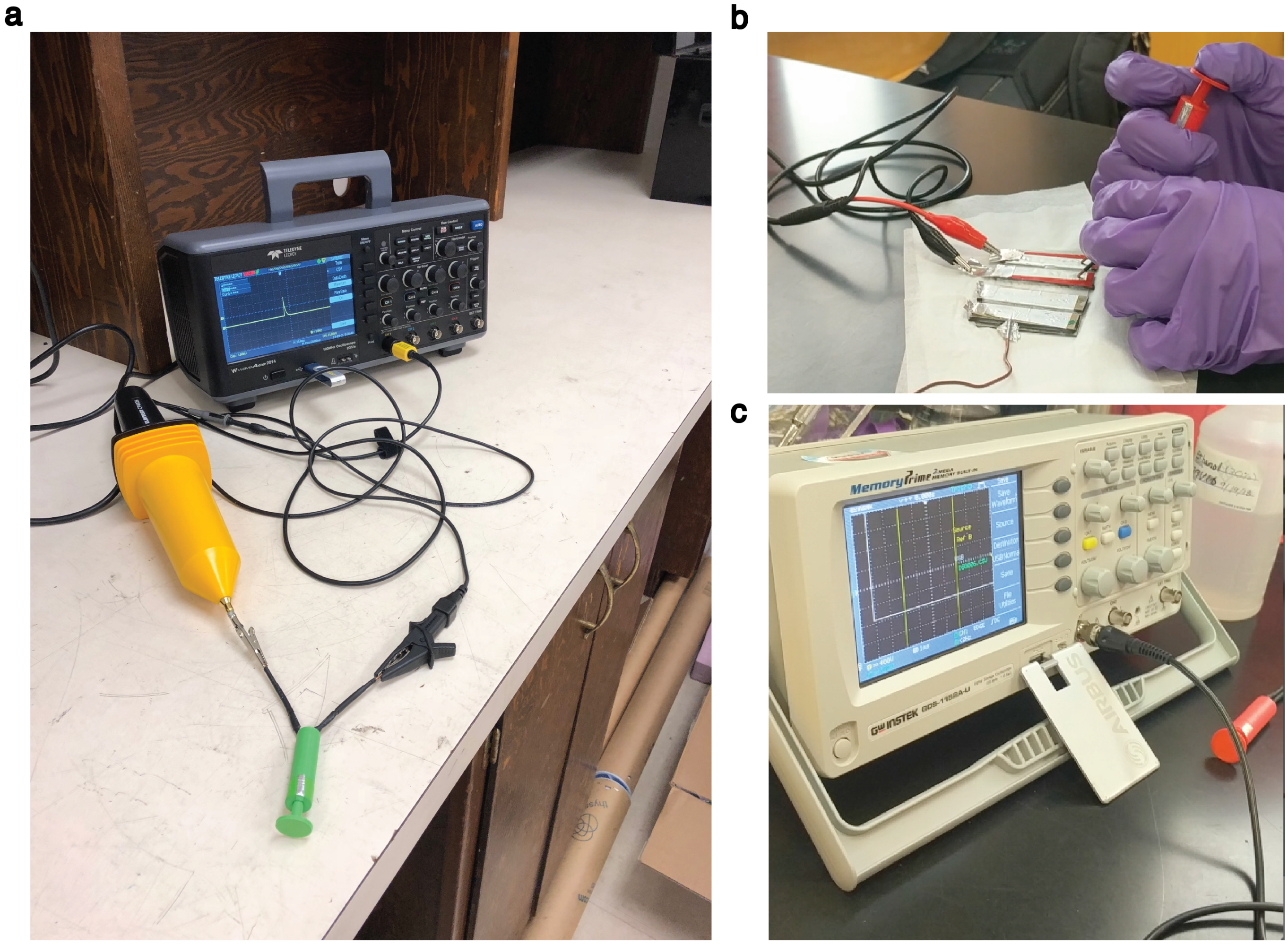


FIG. S5. Setup utilized to quantify voltage output. **a** The oscilloscope was connected to a high-voltage probe along with the ElectroPen. **b** Oscilloscope connections to the cuvette while running the trials to ensure voltage passed through cell suspension. **c** Sample waveform obtained from oscilloscope during trial (voltage clipping is present as ElectroPen voltage output exceeds capacity of the oscilloscope)

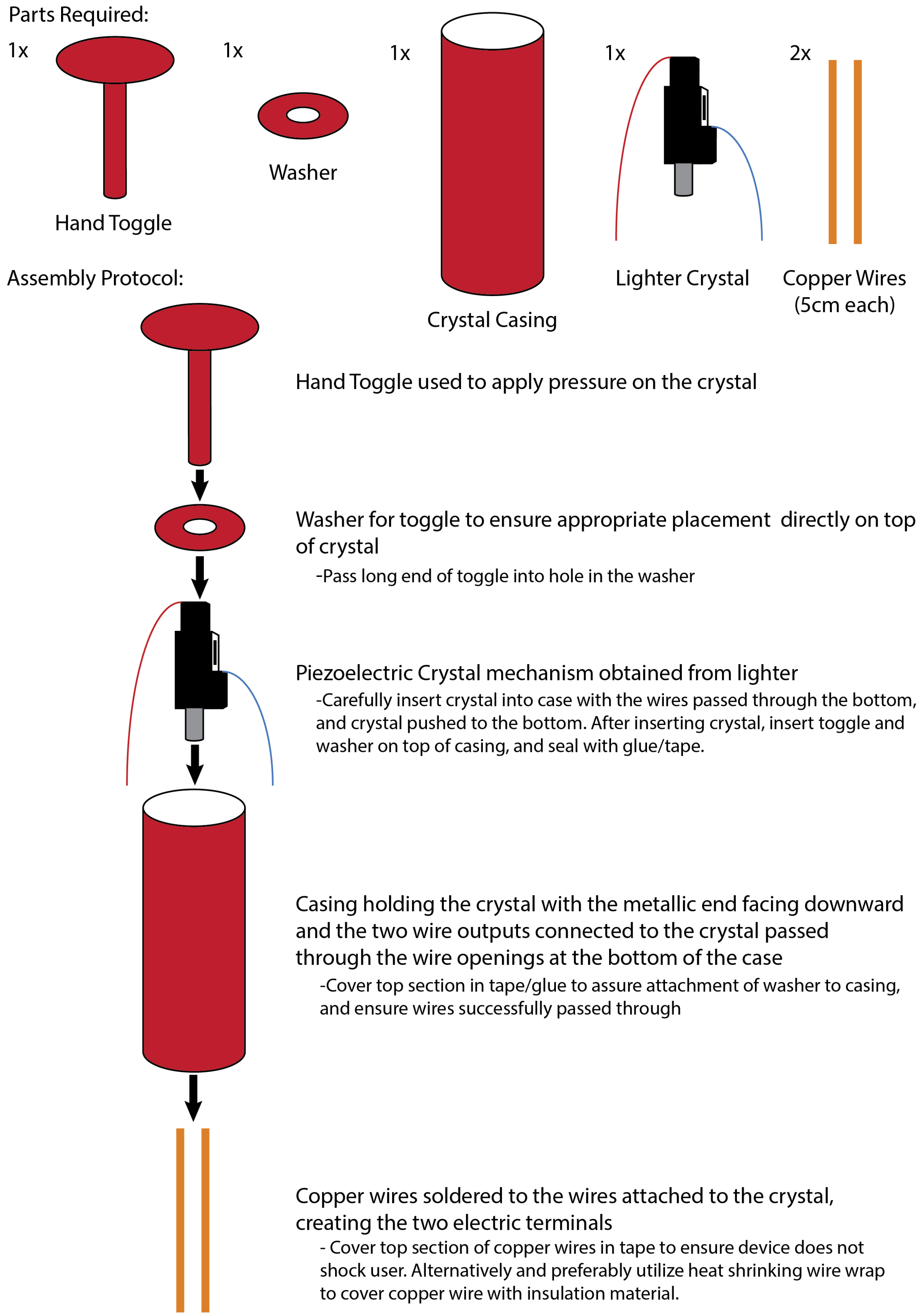


FIG. S6. Illustration of the assembly of the ElectroPen. This depiction indicates the overall construction process with a tutorial found in Supplementary Video S2.

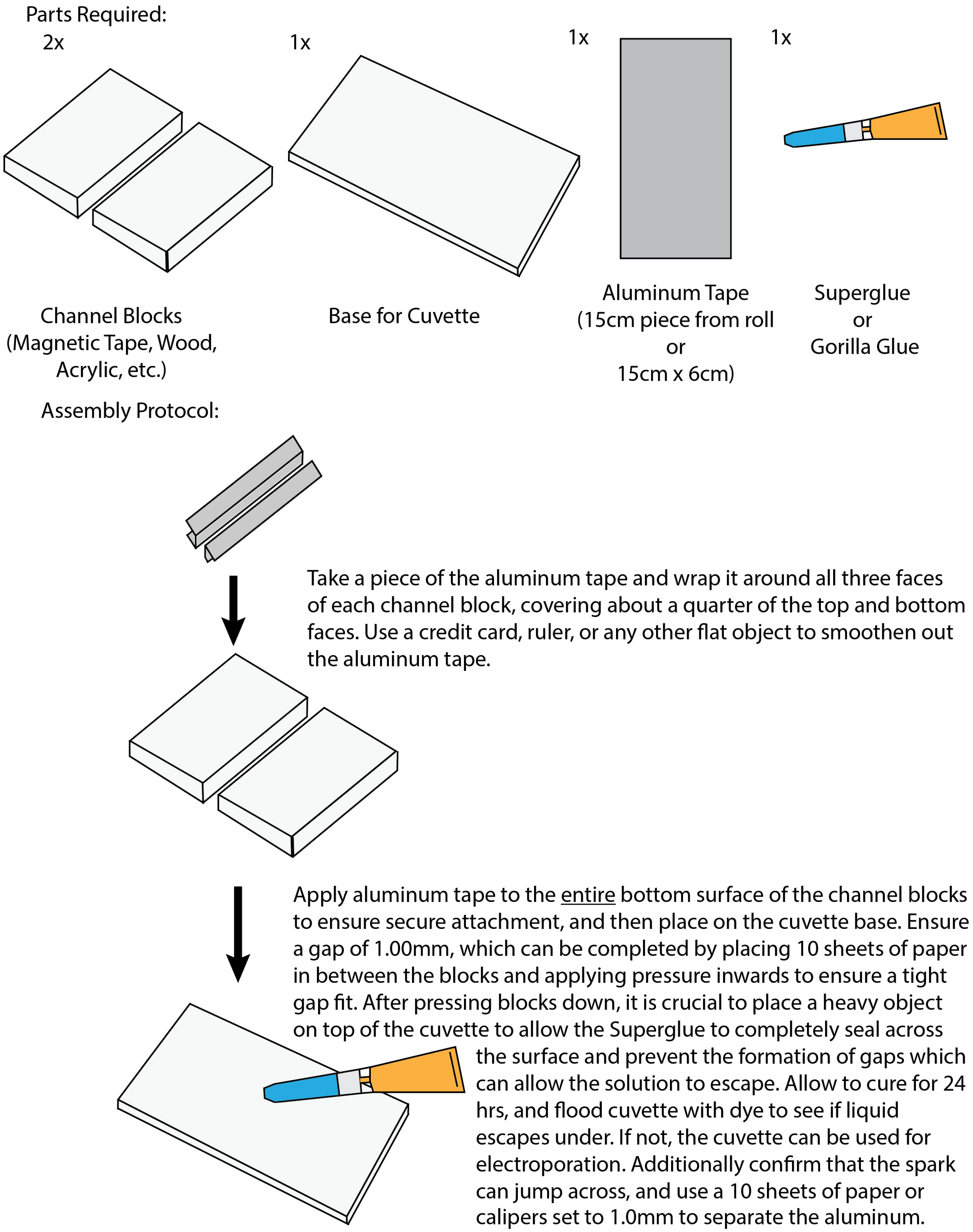


FIG. S7. Illustration of the assembly of the ElectroPen cuvette. This depiction indicates the overall construction process with a tutorial found in Supplementary Video S2.

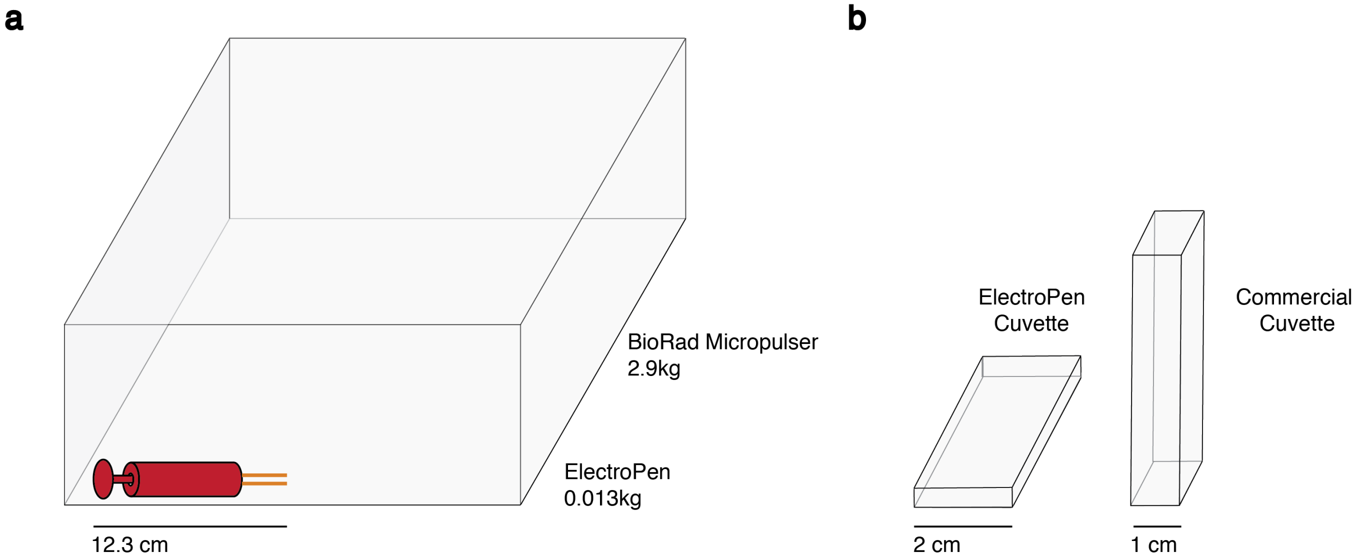


FIG. S8. Size comparison between the ElectroPen and commercial electroporator. **a** Difference in size and weight between the ElectroPen and BioRad Micropulser. **b** Difference in size between commercial electroporation cuvette and ElectroPen cuvette built using a glass slide.

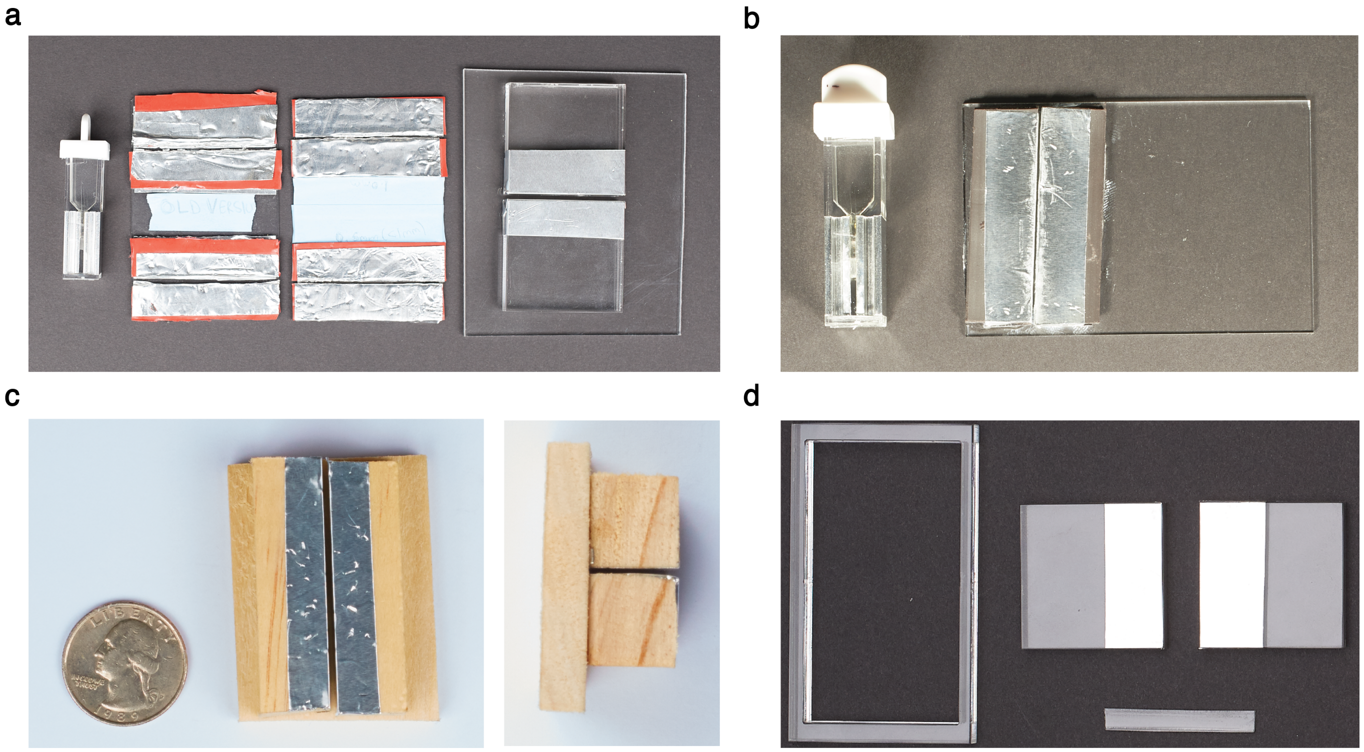


FIG. S9. Depiction of the different variants of the cuvette built using the described basic principles. **a** Pictured left to right is the electroporation cuvette, glass slide cuvettes, and acrylic cuvette. **b** Image of the cuvette used to run the majority of trials in comparison with the commercial cuvette. **c** Cuvette built using wooden blocks. **d** Parts of the acrylic block, with the left piece being the extra surrounding material following laser cutting.

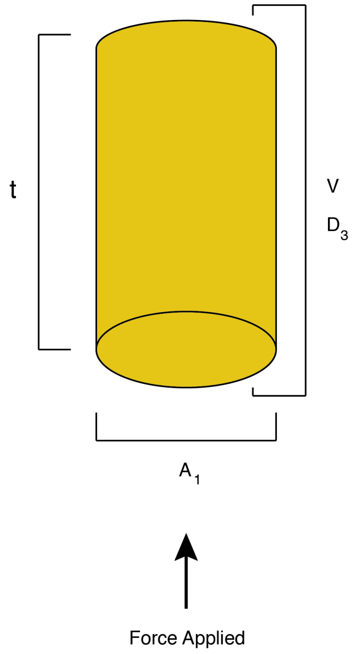


FIG. S10. Depiction of the PZT crystal and the dimensions involved for theoretical model

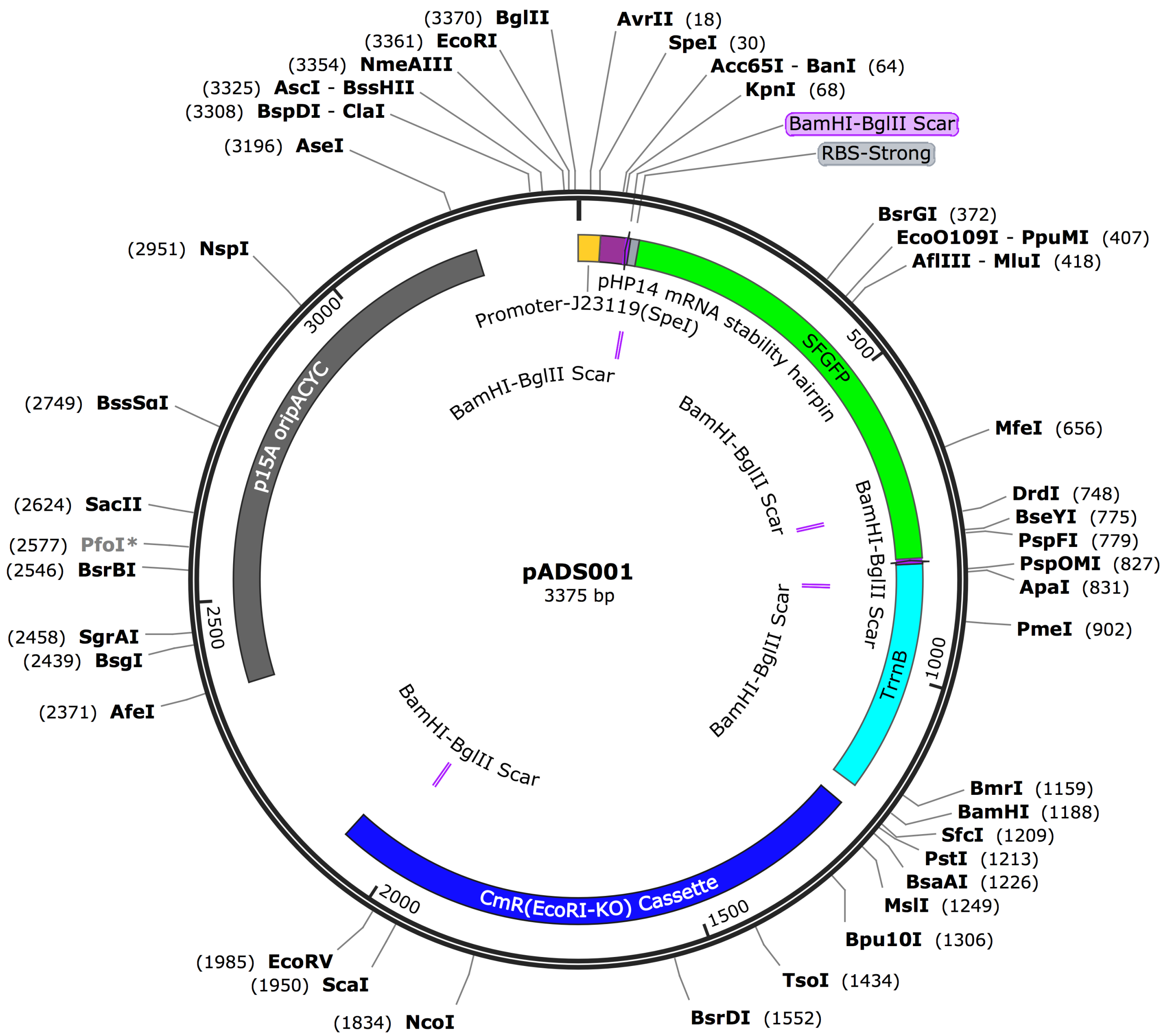


FIG. S11. Diagram depicting the plasmid map of the pADS001 plasmid utilized in the trials conducted. This correlates with the data portrayed in main text Fig. 4(a,b).

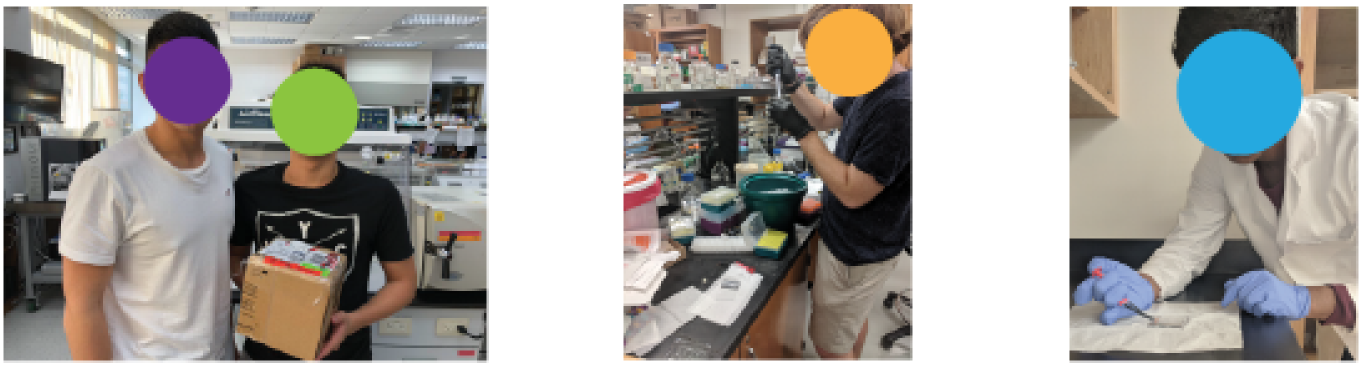


FIG. S12. Images of students from different iGEM Teams across the world testing the ElectroPen. **Authors Note:** Faces of students covered in accordance with **bioRxiv** policy to ‘avoid the inclusion of photographs and any other identifying information of people because verification of their consent is incompatible with the rapid and automated nature of preprint posting’.
